# Building customizable auto-luminescent luciferase-based reporters in plants

**DOI:** 10.1101/809533

**Authors:** Arjun Khakhar, Colby Starker, James Chamness, Nayoung Lee, Sydney Stokke, Cecily Wang, Ryan Swanson, Furva Rizvi, Takato Imaizumi, Dan Voytas

## Abstract

Bioluminescence is a powerful biological signal that scientists have repurposed to design reporters for gene expression in plants and animals. However, there are some downsides associated with the need to provide a substrate to these reporters, such as its high cost and non-uniform tissue penetration. In this work we reconstitute a fungal bioluminescence pathway (FBP) *in planta* using an easily composable toolbox of parts. We demonstrate that the FBP can create luminescence across various tissues in a broad range of plants without external substrate addition. We also show how our toolbox can be used to deploy the FBP *in planta* to build auto-luminescent reporters for the study of gene-expression and hormone fluxes. A low-cost imaging platform for gene expression profiling is also described. These experiments lay the groundwork for the future construction of programmable auto-luminescent plant traits, such as creating light driven plant-pollinator interactions or light emitting plant-based sensors.

## Introduction

Bioluminescence is one of nature’s more spectacular tricks. It is used by a diverse set of organisms to achieve a broad range of goals such as attracting mates, scaring off predators and recruiting other creatures to spread spores^1,2,3,4,5^. The mechanism of light emission is broadly conserved: an enzymatic oxidation reaction by a luciferase enzyme turns a luciferin substrate into a high energy intermediate which decays to produce light^1^. There have been several different examples of luciferin substrate-luciferase enzyme pairs described to date^6,7^. Researchers have leveraged some, such as the firefly and renilla luciferase enzymes to build reporters to study gene expression in plants and other eukaryotic systems^8,9^. These reporters have a high signal to noise ratio because there is effectively no background signal produced by plants.

Additionally, thanks to its water-soluble luciferin substrate, firefly luciferase can be used to visualize live changes in gene expression in plants in a non-invasive manner, theoretically enabling whole plant time lapse imaging^8^. However, current bioluminescent reporters do present some significant challenges which has limited their broad application for macro scale visualization of gene expression. One major challenge is uniform delivery and penetration of the luciferin substrate, especially in adult plant tissues. Some luciferins like coelenterazine, the substrate for renilla luciferase, are non-water soluble and so cannot be used for imaging without cell-lysis^9^. The luciferin for firefly luciferase, D-luciferin, is water soluble and can be topically applied to whole plants or delivered through watering^10^. However, uniform substrate delivery is challenging to achieve, which makes it difficult to disambiguate whether a lack of bioluminescent signal is due to a lack of expression of the luciferase or delivery of the substrate. Additionally, the relatively high cost of D-luciferin (up to several hundred dollars per gram) makes continuous whole plant imaging a prohibitively expensive practice.

To avoid the need to deliver the luciferin substrate, an optimal solution would be to engineer plants such that the substrate is produced in every cell. Krichevsky et al. demonstrated one approach to achieve this by incorporating the lux operon from the prokaryote *Photobacterium leiognathi* into the chloroplast genome in *Nicotiana tabacum*^11^. While auto-luminescence was achieved, this approach is extremely challenging to extend to other plants due to the technical challenges of transforming plastids. In their 2018 paper, Kotlobay and Sarkisyan et al. identify a fungal bioluminescence pathway (FBP) consisting of a set of three genes that are sufficient to convert caffeic acid, a common plant metabolite^12^, into a luciferin molecule and a paired luciferase, *Luz*, that can oxidize it to create a bioluminescent signal^12^. This pathway can be nuclear encoded, as it is derived from a eukaryotic fungus, overcoming the challenges associated with plastid transformation. Additionally, this luciferase produces light in the green spectrum, which minimizes absorption by the predominant plant pigment, chlorophyll, and makes it spectrally separable from the other major luciferase-based reporters^13^.

In this report we sought to reconstitute the FBP *in planta* and to test if natively produced caffeic acid could be used to generate bioluminescence. We also tested whether the FBP functioned across plant species and if it could be used to study the spatiotemporal patterns of gene expression. In this work we describe a toolkit of resources to easily generate FBP-based reporters to study gene expression *in planta*. We also demonstrate how this toolkit could be deployed to generate programable auto-luminescence patterns *in planta* and to build plant-based biosensors with luminescent outputs that do not require external substrate addition.

## Results

### Reconstitution of the fungal bioluminescence pathway *in planta* leads to auto-luminescence

To test if the fungal bioluminescence pathway described by Kotlobay and Sarkisyan et al.^12^ would function *in planta*, we attempted to transiently reconstitute this pathway in the leaves of *Nicotiana benthamiana*. We chose to use *N. benthamiana* because Agrobacterium infiltration provides an easy way to transiently express genes. We synthetized codon optimized versions of the three genes necessary to turn caffeic acid into 3-hydroxyhispidin, 4′-phosphopantetheinyl transferase (*NPGA*) from *Aspergillus nidulans*, hispidin-3-hydroxylase (*H3H*) from *Neonothopanus nambi*, and a polyketide synthase (*Hisps*) also from *N. nambi*^12^ (Figure 1A). We also generated a codon optimized version of the fungal luciferase, *Luz*. These DNA sequences were also domesticated to remove all commonly used type II-S restriction sites to make them compatible with the MoClo plasmid assembly kit^14^. This enabled the promoters and terminators to be easily swapped and facilitated the creation of a toolbox of vectors to build FBPs for constitutive or spatio-temporally regulated auto-luminescence (Table 1). The expression cassettes can be easily assembled into single T-DNAs that express the entire FBP with a one-step GoldenGate reaction.

**Table 1.**
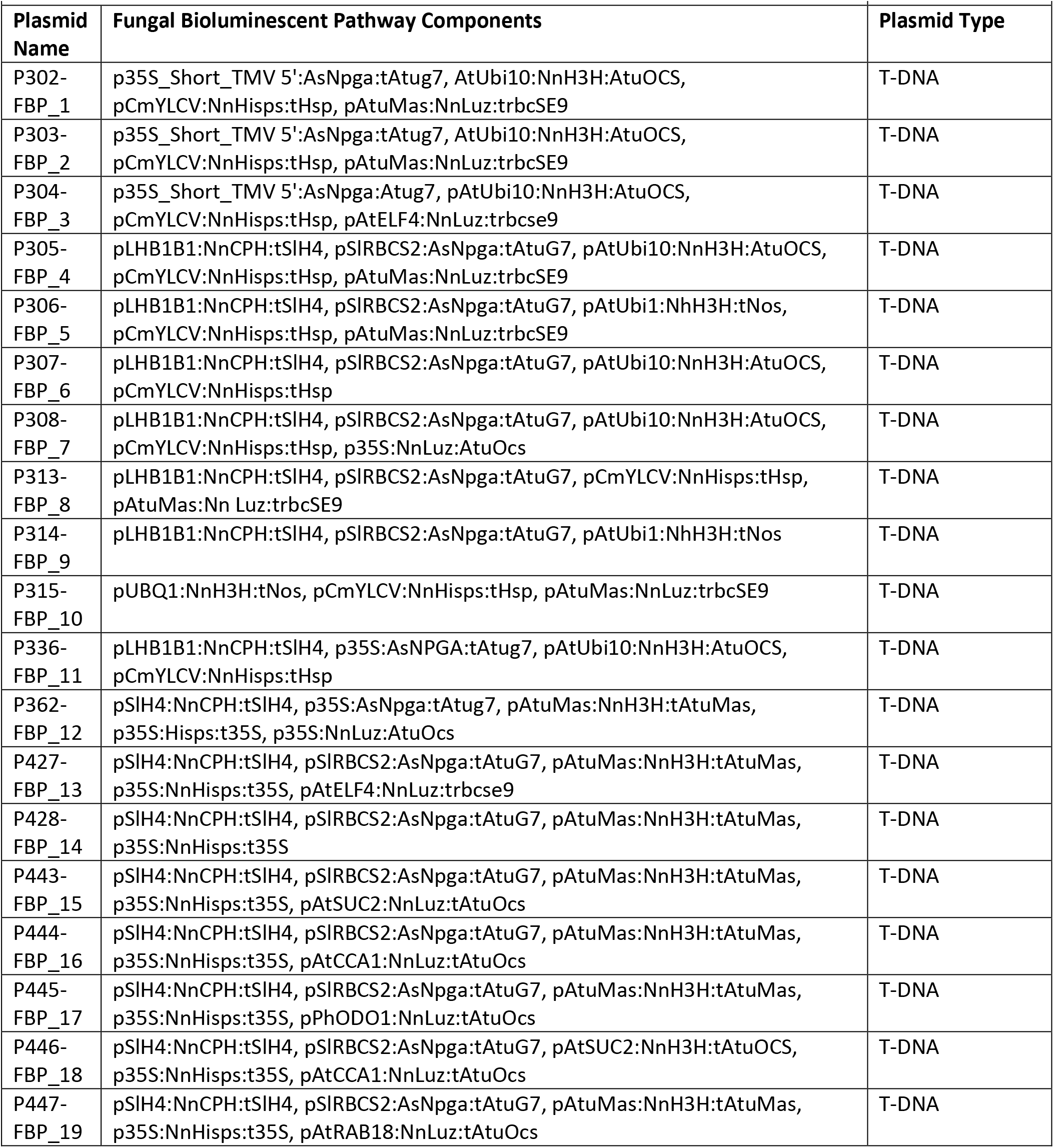
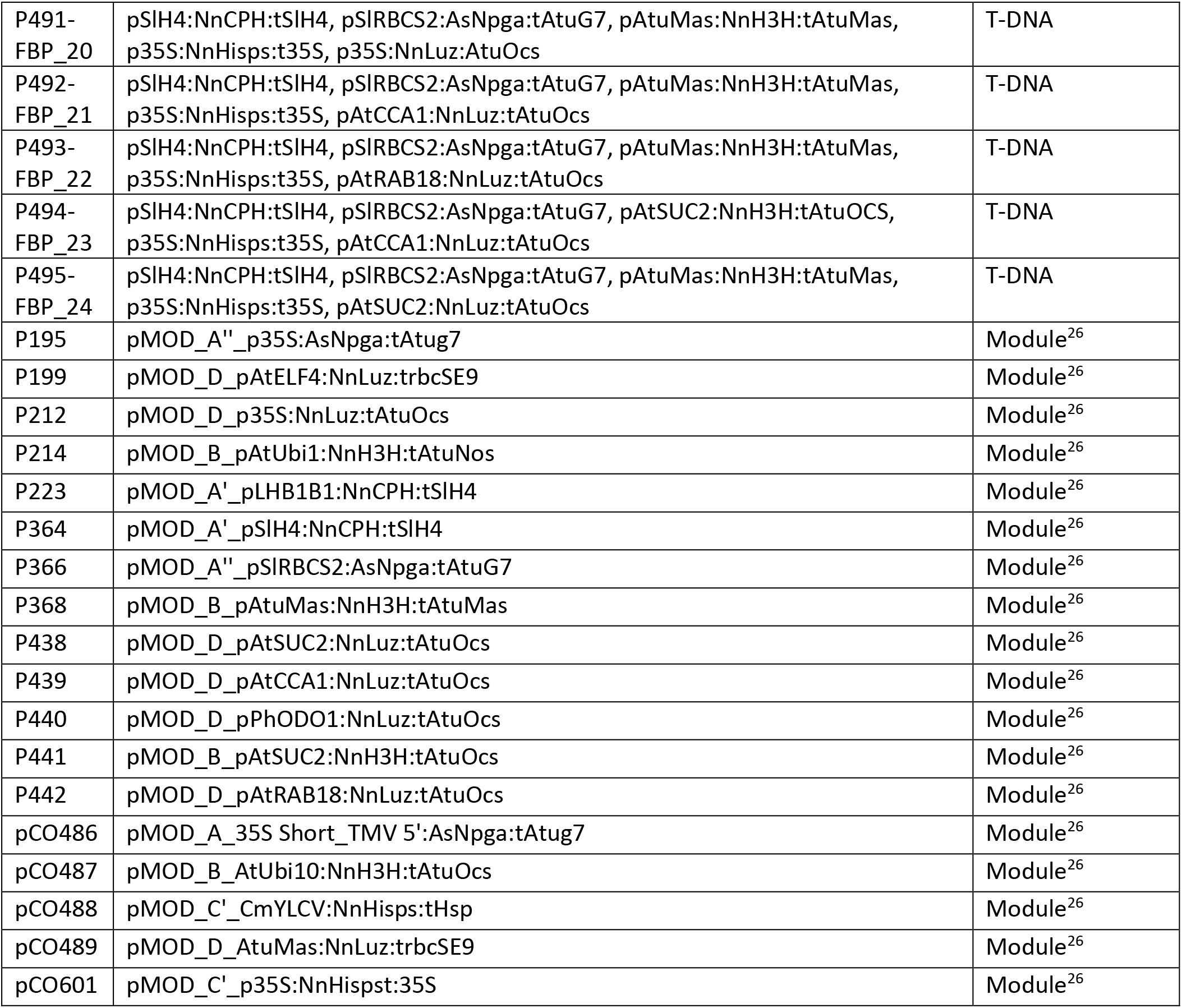
Vectors containing FBP components.

**Figure 1.**
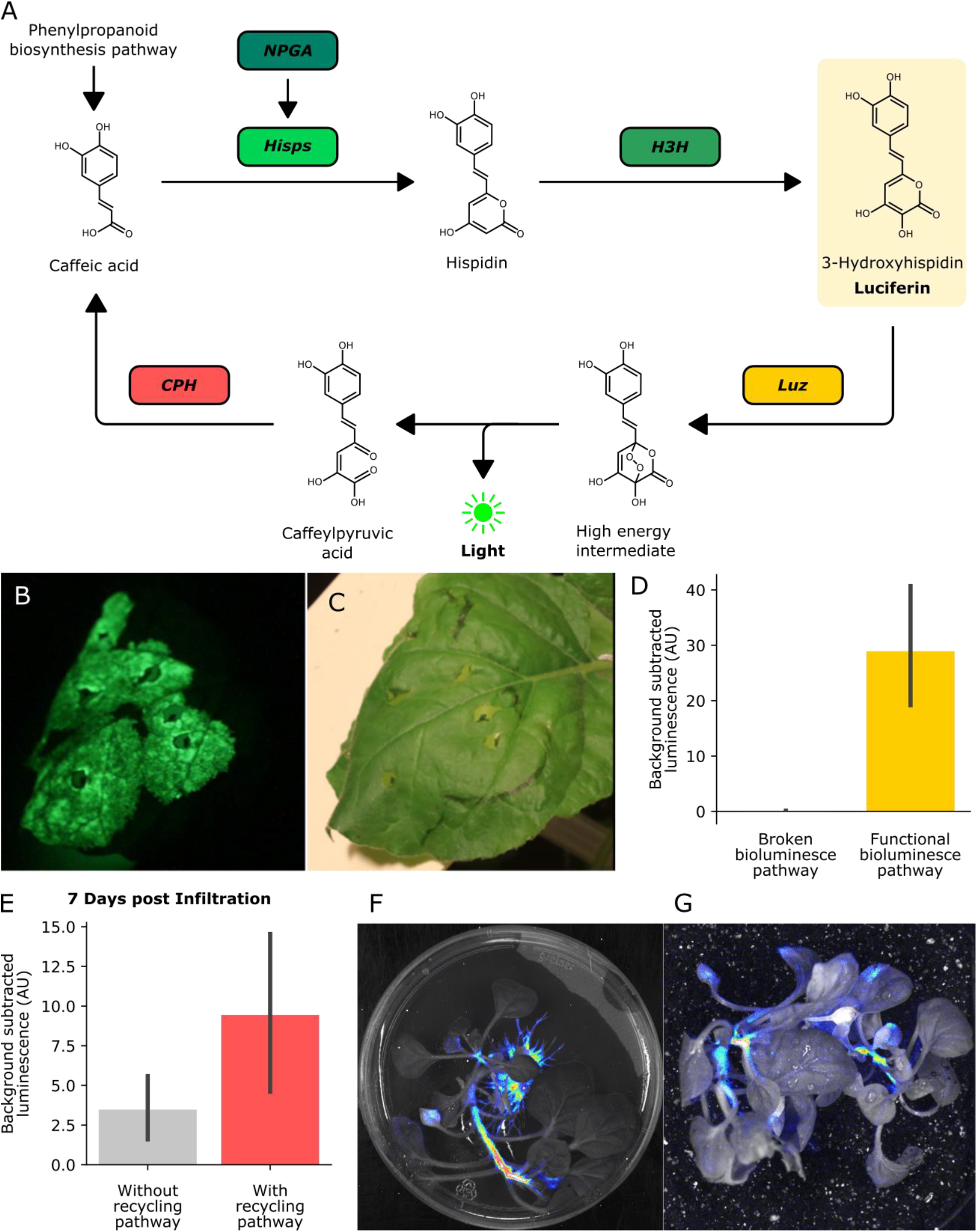
A) Schematic of the chemical reactions driving the generation of auto-luminescence *in planta*. Caffeic acid produced by the phenylpropanoid biosynthesis pathway is converted to Hispidin by *Hisps* once it is post-translationally activated by *NPGA*. Hispidin is then converted to 3-Hydroxyhispidin, the luciferin molecule, by *H3H*. Finally, the luciferase *Luz* oxidizes 3-Hydroxyhispidin to a high energy intermediate which degrades into Caffeylpyruvic acid, producing light. *CPH* can turn Caffeylpyruvic acid back into Caffeic acid, closing the cycle. B) An image in the dark with an eight minute exposure of a *N. benthamiana* leaf infiltrated with the FBP demonstrating auto-luminescence in the infiltrated zone. C) An image in the light of the same leaf with the infiltrated zone marked with a black outline. D) Bar plots visualizing background subtracted luminescence from *N. benthamiana* leaves infiltrated with either a functional FBP (yellow) or a broken control (gray) missing the luciferase, *Luz*, three days after infiltration (n=12, p=0.0002). Black bars represent standard deviation. E) Bar plots visualizing background subtracted luminescence seven days after infiltration from *N. Benthamiana* leaves infiltrated with FBPs that either have (pink) or do not have (gray) the *CPH*-based recycling pathway (n=12, p=0.05). Black bars represent standard deviation. F, G) Luminescence signal captured with a CCD camera superimposed on a bright field image of a transgenic *N. benthamiana* plant with a genomically integrated FBP on rooting media (F) and in soil (G). Warmer colors correspond to higher luminescence.

When we imaged *N. benthamiana* leaves three days after delivery of the FBP via Agrobacterium infiltration, we see significant bioluminescence over background at the site of infiltration (Figure 1B,C,D), demonstrating the pathway can produce bioluminescence using natively synthesized caffeic acid. Upon comparing luminescence from leaves that had the complete pathway versus controls in which enzymes from the pathway were missing, we only observe a bioluminescence signal with the complete pathway (Figure 1D, Supplement figure 1). This indicates that there are no native enzymes expressed at functional levels in *N. benthamiana* leaves which could complement the functions of the FBP enzymes. We also observe no bioluminescence from confluent cultures of Agrobacterium that carry the pathway (Supplementary figure 1), implying the observed bioluminescent signal is not from expression of this pathway in Agrobacterium.

### Incorporation of spent luciferin recycling pathway prolongs transient auto-luminescence

In their paper, Kotlobay and Sarkisyan et al^12^ also describe an additional enzyme, caffeylpyruvate hydrolase (*CPH*), that recycles the spent luciferin, caffeylpyruvic acid, back into the substrate for the pathway, caffeic acid (Figure 1A). We reasoned that incorporating this enzyme into our reconstituted pathway would create a stronger bioluminescence signal due to a higher substrate concentration and potentially prolong the presence of transient bioluminescence from Agrobacterium infiltration. To test this hypothesis, we built a new pathway with all the enzymes described above as well as one additional expression cassette for *CPH* expression. When we compared luminescence from Agrobacterium infiltrated *N. benthamiana* leaves, we observed approximately equal signals at four days, consistent with initial saturating substrate conditions (Supplementary Figure 2). At later time points we see higher mean luminescence signal from leaves infiltrated with the *CPH* containing pathway, consistent with the hypothesis that when the substrate is limited, the presence of a recycling pathway improves the bioluminescence signal (Figure 1E). Based on these results, for all further experiments we used FBPs that included the recycling enzyme, *CPH*.

### Stable integration of the fungal bioluminescence pathway creates auto-luminescent plants

Our experiments thus far implied that we could create auto-luminescent plants by genomically integrating expression cassettes for the three enzymes in the biosynthesis pathway, namely *NPGA*, *H3H*, and *Hisps*, the recycling pathway, *CPH*, and the fungal luciferase, *Luz*. We used the FBP characterized in Figure 1 to generate a stable transgenic line of *N. benthamiana*. When we characterized the luminescence on rooting medium (Figure 1F) and in soil (Figure 1G), we observed auto-luminescence across the plant. We see stronger signals in the root tips and shoot apical meristem, consistent with a higher density of cells in these tissues. Similar patterns of expression are observed in root tips of *Arabidopsis thaliana* with constitutively expressed firefly luciferase^8^. It is therefore possible to generate plants with genomically integrated FPBs to create stable auto-luminescence.

### Auto-luminescence in different plant species

While *N. benthamiana* is an excellent model to prototype the FBP, we hypothesized that it should, in principle, work in any plant capable of producing caffeic acid, as long as the appropriate promoters and terminators were used to ensure expression of the pathway enzymes. To test this hypothesis, we performed transient Agrobacterium-based delivery of the FBP into the model plant *A. thaliana* and the crop plant *Solanum lycopersicum* (tomato) using the AgroBEST protocol^15^. We observed robust auto-luminescence signal over background in the cotyledons of treated seedlings (Figure 2A,B). We also performed Agrobacterium infiltrations of the leaves of the ornamental plant, Dahlia, and were able to observe auto-luminescence (Figure 2C). Finally, we explored if the FBP was functional in petals, as luminescence of this tissue has applications for horticulture as well as engineering plant pollinator interactions. Agrobacterium infiltrations with the FBP of petals from *Catharanthus roseus* (periwinkle), *Petunia axillaris*, and three different varieties of *Rosa rubiginosa* (Rose) resulted auto-luminescence in all flowers with a range of intensities (Figure 2D,E, Figure 3A). We also observed that the signal dropped off within a day of the flowers being detached from the plant body, whereas it persisted when the flowers remained connected to the plant body. This might mean some of the pathway substrates or co-factors might be trafficked from source tissues; however, more tests are required to confirm this hypothesis. These results demonstrate that the FBP can function across a wide range of plants.

**Figure 2.**
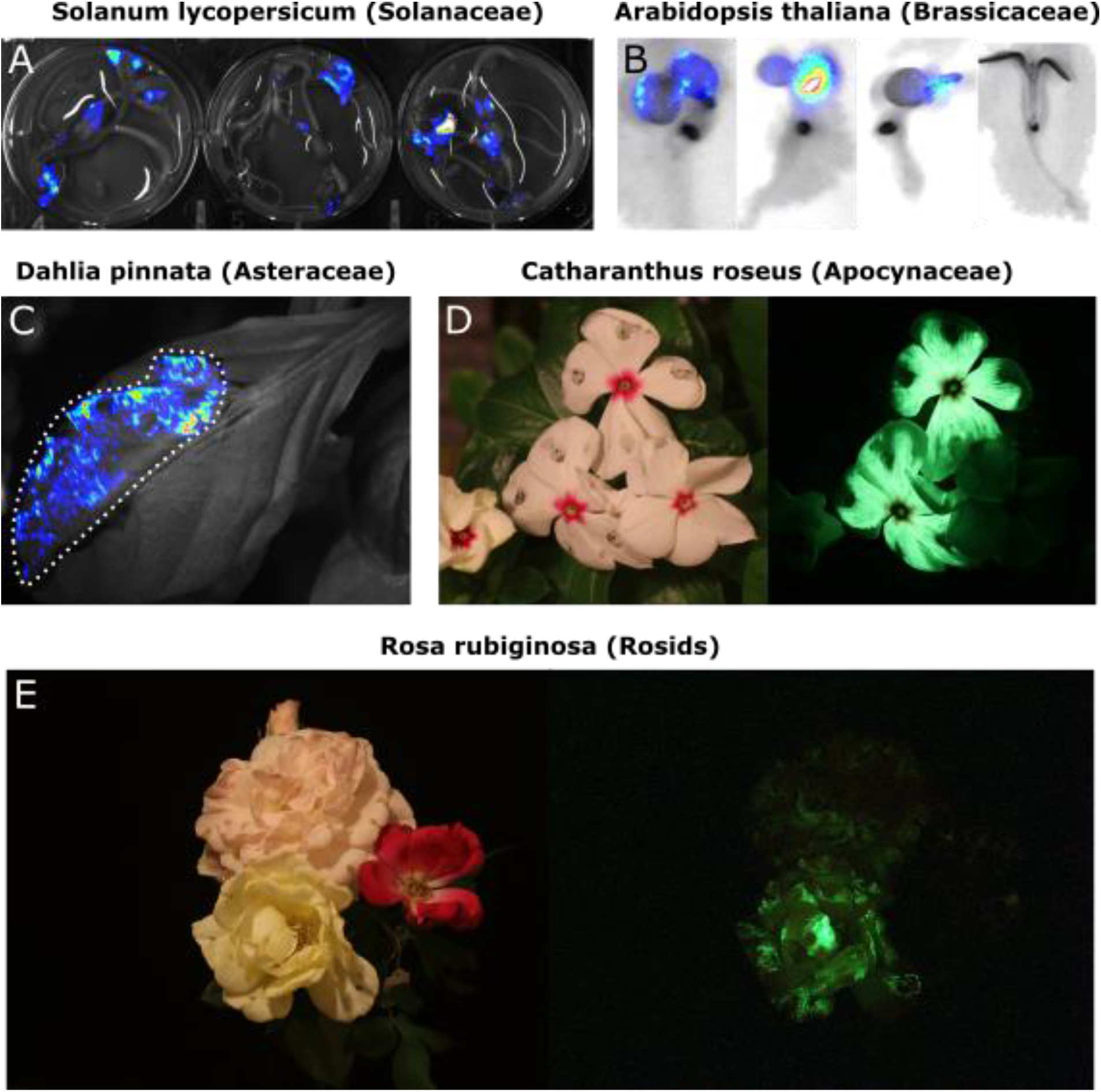
A) Luminescence signal superimposed on a bright field image of *Solanum lycopersium* (tomato) seedlings with an FBP delivered to them via the agrobest protocol. Warmer colors correspond to higher luminescence signal. B) Luminescence signal superimposed on bright field images of *Arabidopsis thaliana* seedlings with a functional FBP (left three seedlings) or a broken FBP (right most seedling) delivered to them via the agrobest protocol. Warmer colors correspond to higher luminescence signal. C) Luminescence signal superimposed on a bright field image of a *Dahlia pinnata* leaf infiltrated with an FBP. D) Lit (left) and unlit, long exposure (right) true color images of *Catharanthus roseus* flowers infiltrated with an FBP. E) Lit (left) and unlit, long exposure (right) true color images of *Rosa rubiginosa* flowers infiltrated with an FBP.

**Figure 3.**
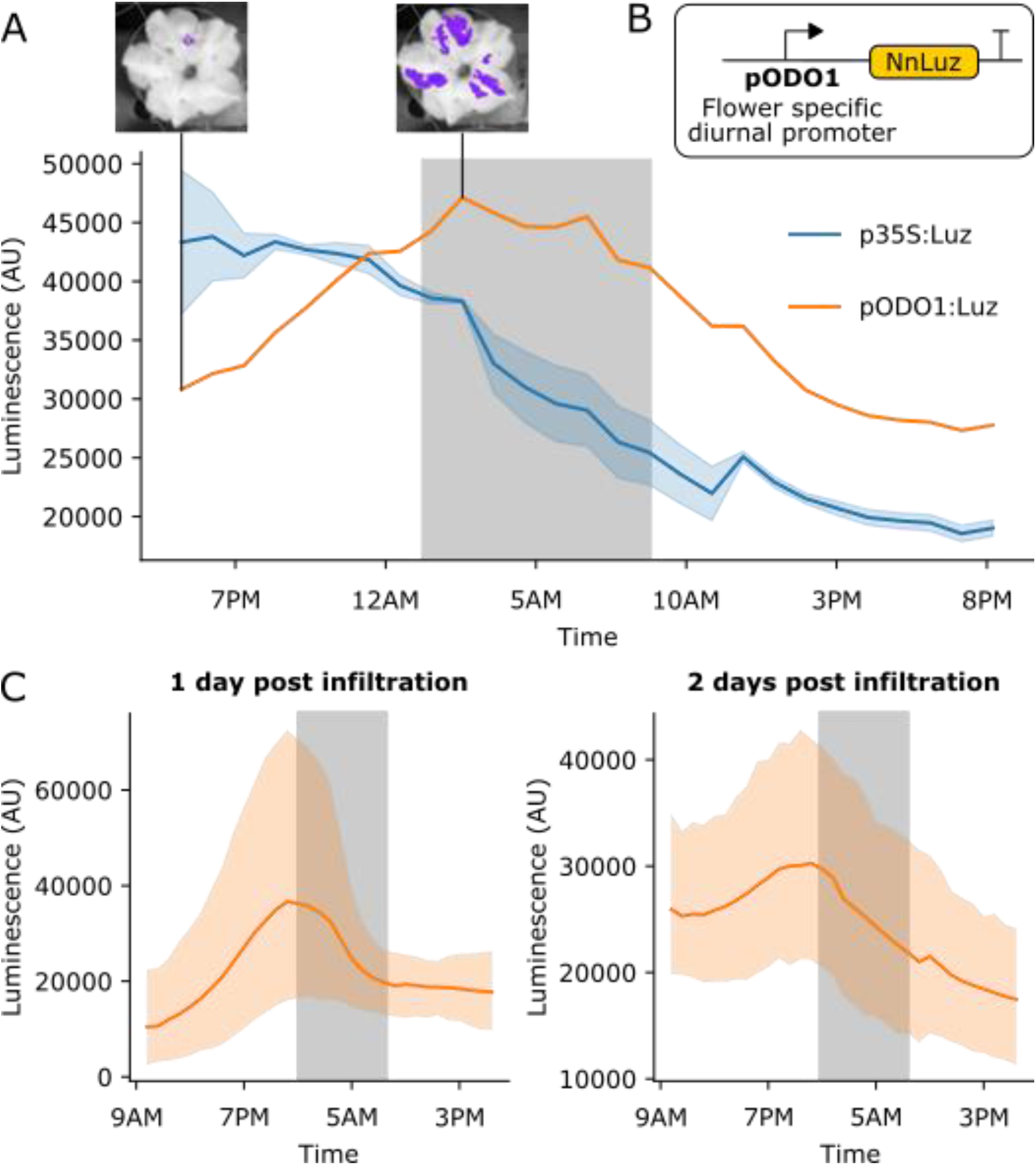
A) Time course luminescence data from long day entrained *P. axillaris* flowers that were infiltrated with FBPs. The orange line represents FBP with *Luz* driven by the *ODO1* promoter from Petunia (n=1) and the blue line represents the mean luminescence signal of an FBP with *Luz* driven by the 35S promoter (n=2). The light blue outline represents standard deviation. The gray background represents the night period. Representative images of petunia flowers infiltrated with the pODO1:*Luz* at minimum and maximum bioluminescence are also shown, where luminescence signal is overlaid on the bright field image. B) A schematic of the *Luz* expression cassette being driven by flower specific diurnal promoter of the petunia gene, *ODO1*, which is expected to have peak expression at the day to night transition. C) Time course luminescence data from long day entrained *P. axillaris* flowers that were infiltrated with an FBP with *Luz* driven by the *ODO1* promoter, 1 and 2 days post infiltration (n=7 for both). The solid orange lines are mean luminescence and light orange outlines represent standard deviation. The gray background represents the night period.

### Visualizing spatio-temporal patterns of gene expression

*In planta* substrate-independent bioluminescence promises to be a powerful reporter technology, as it enables long-term characterization of gene expression on a macro scale in a cost-effective manner. The approach also does not suffer from the issues associated with current bioluminescent reporters, such as the challenges of substrate application and non-uniform substrate penetration. To validate that the pathway could be used to study spatio-temporal patterns of gene expression, we built a version of the pathway in which the expression of *Luz* was driven by the promoter of the *ODORANT1* (*ODO1*) gene from petunia^16,17,18^ (Figure 3B). This gene has been previously characterized with a firefly luciferase-based reporter. Its expression was only observed in the flowers of petunia and was shown to be diurnal, peaking in the evening at the transition of day to night^16^. We used Agrobacterium infiltration to deliver two versions of FBP to petunia flower petals: one with *Luz* expressed from the *ODO1* promoter (pODO1) and the other with *Luz* expressed from the constitutive 35S promoter. We then performed time course imaging of detached flowers. We observed an increase in luminescence in the evening at the transition between day and night in the flowers treated with the pODO1*:Luz*, a trend not seen in the 35S*:Luz* control (Figure 3A).

Visualizing multiple diurnal oscillations in flowers is technically challenging, as the auto-luminescence signal only lasts approximately one day after the flower is separated from the plant body, a necessity for mounting it in the imaging platform. To overcome this limitation, we infiltrated two sets of flowers 24 hours apart and then harvested tissue and began time course imaging on the same day. In this way we were able to quantify luminescence signal from the pODO1*:Luz* over a two-day period post infiltration. We observed the expected luminescence signal peak at the transition between day and night both one and two days after infiltration (Figure 3C), with a diminishing signal over time, which can be explained by the gradual loss of T-DNA expression over time^19^ combined with gradual senescence of the detached flower. These results demonstrate that the FBP can be used in a similar fashion to firefly luciferase to study spatio-temporal patterns of gene expression without necessitating substrate addition. These experiments also serve as a proof-of-concept that, by using an appropriate set of promoters to drive the pathway genes, an FBP could be programmed to generate specific spatio-temporal patterns of auto-luminescence in stable transgenic lines for synthetic biology applications.

### Visualizing hormone fluxes *in planta*

Hormone signals coordinate diverse aspects of plant metabolism and development by carrying information from cell to cell across tissues and organ systems in the plant. Firefly luciferase-based reporters have been an invaluable tool to study how hormone fluxes can trigger developmental changes in the plant^8^. However, the challenges of substrate cost, application and penetration make using them across the entire plant body or over long periods problematic. The substrate independence of the FBP makes it an attractive alternative to overcome these challenges. To demonstrate how this pathway could be engineered to build a hormone biosensor we built a version of the pathway with the *AtRAB18* promoter (pRAB18) driving expression of *Luz* (Figure 4A). This promoter has been previously characterized as having strong and selective increase in expression in response to the plant hormone abscisic acid (ABA)^20^.

**Figure 4.**
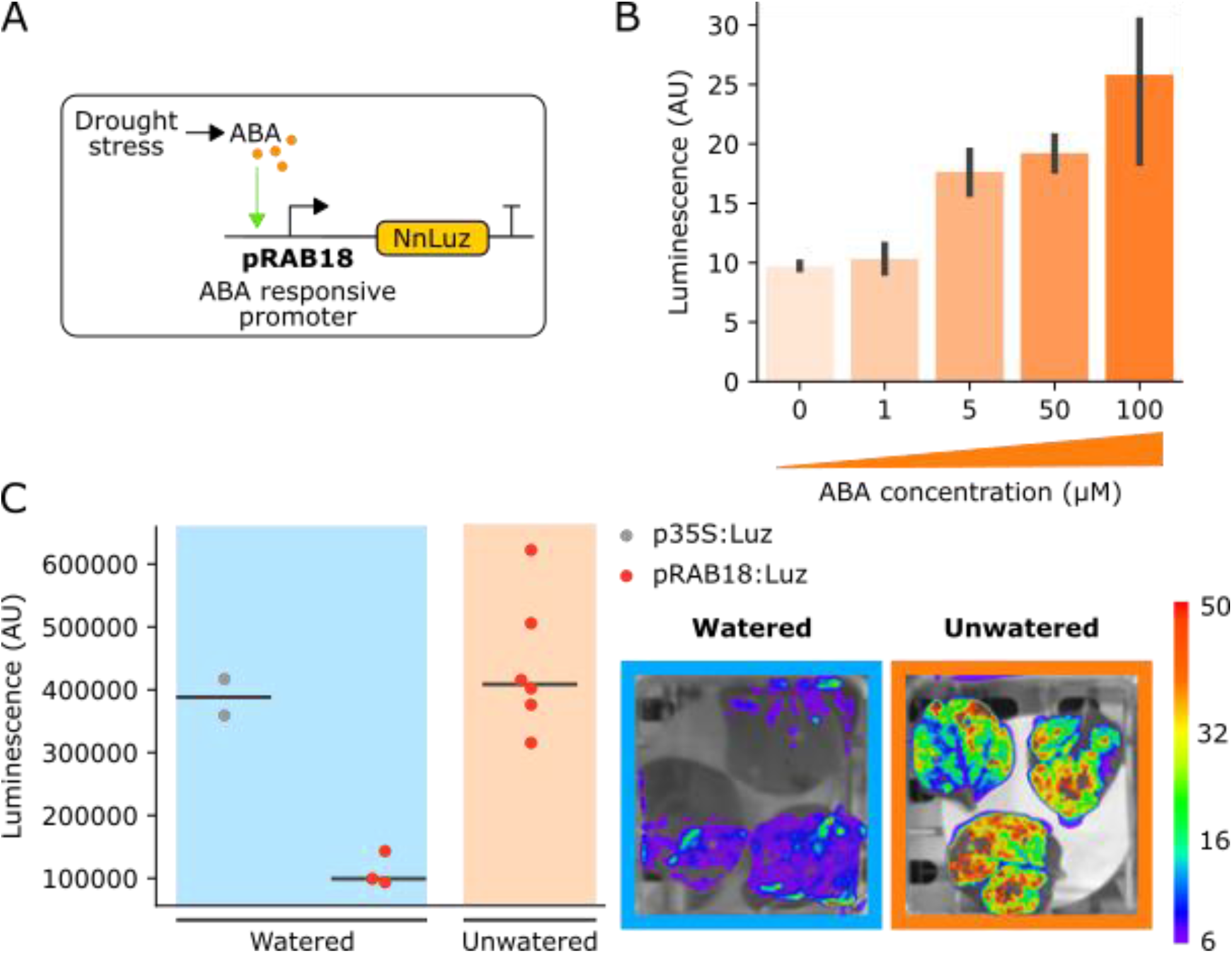
A) Schematic of the *Luz* expression cassette driven by the *AtRAB18* promoter which responds to ABA signals, such as those that produced in response to drought. B) Bar plots summarizing mean luminescence signal observed from *N. benthamaina* leaves that were co-infiltrated with the pRAB18:*Luz* containing FBP along with increasing concentrations of the hormone ABA and then imaged after three days. Black bars represent standard deviation (n=3). C) Luminescence signal observed from *N. benthamaina* leaves that were infiltrated with an FBP and then were either kept moist (blue background) or were allowed to desiccate to trigger an ABA signal (orange background). Gray and red dots represent data from independent leaves infiltrated with FBP with a 35S or pRAB18 driven *Luz* respectively. Representative images of watered and un-watered leaves infiltrated with the pRAB18:*Luz* FBP with luminescence signal overlaid on a bright field picture. Warmer colors represent higher signals.

We first validated if this FBP could indeed faithfully report ABA levels *in planta*. To do this, we assayed the bioluminescent signal produced three days after infiltration into *N. benthamiana* leaves that were treated with increasing concentrations of ABA. As expected, we observed increasing bioluminescence signal with increasing doses of ABA (Figure 4B).

We next tested if this system could respond to endogenous ABA signals. The pRAB18 driven FBP was delivered to the leaves of *N. benthamiana* via Agrobacterium infiltration. One set of leaves was allowed to desiccate, while the other set was kept moist. The bioluminescence of the leaves was then imaged over time. We expected that as the leaves dry out, they would begin to produce ABA^21^, leading to an increase in luminescence. We observed a significant increase in luminescence in desiccating leaves, up to levels of an FBP with a 35S:*Luz* (Figure 4C). We did not see an increase in signal in leaves that were kept moist (Figure 4C,D). These results demonstrate the utility of the fungal luciferase as a tool to enable substrate independent visualization of hormone fluxes *in planta*. They also serve as a proof of concept for how this pathway could be deployed in stable plant lines as a biosensor for soil moisture levels in the future.

### Developing a low-cost, long-term bioluminescence imaging system

The power of bioluminescence as a tool to study dynamic signals in biology, such as ABA accumulation during drought, is challenging to access in lower resource settings due to high costs of substrate and instrumentation. The substrate independence of the FBP makes it more economical than other bioluminescent systems. However, the high cost and small chamber size of commercially available luminescence imaging setups make running multiple experiments in parallel cost prohibitive. The cost of such commercial systems can exceed $100,000USD. To address this challenge, we leveraged a wealth of open-source software and off-the-shelf hardware components to design and build a relatively low cost, modular platform for bioluminescence imaging. Our platform uses a Raspberry Pi single board computer as a controller for a DSLR camera, outfitted with a macro lens and mounted in either a dark room or pop-up plant growth tent. The code used for controlling customized still and time-course imaging with this platform, as well as the image processing pipeline, is available on Github (Table 2). The entire cost comes to less than $2,500, the bulk of which is for the camera and lens; substituting an entry-level DSLR and kit lens would bring the cost below $2,000.

**Table 2.**
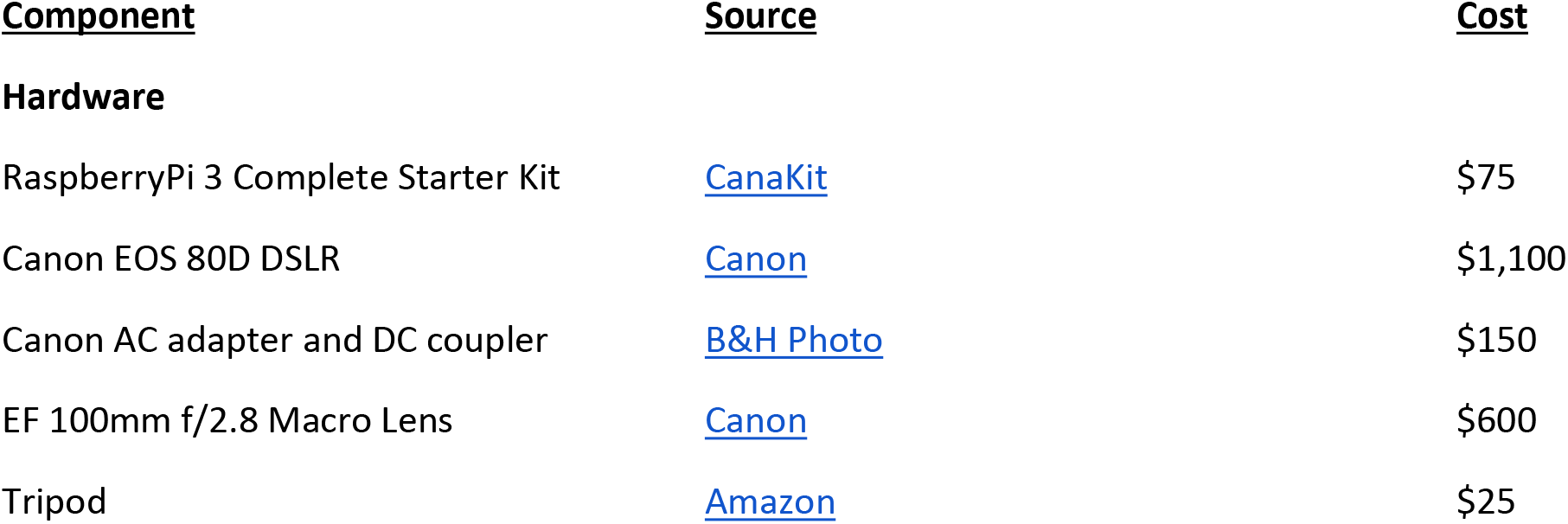
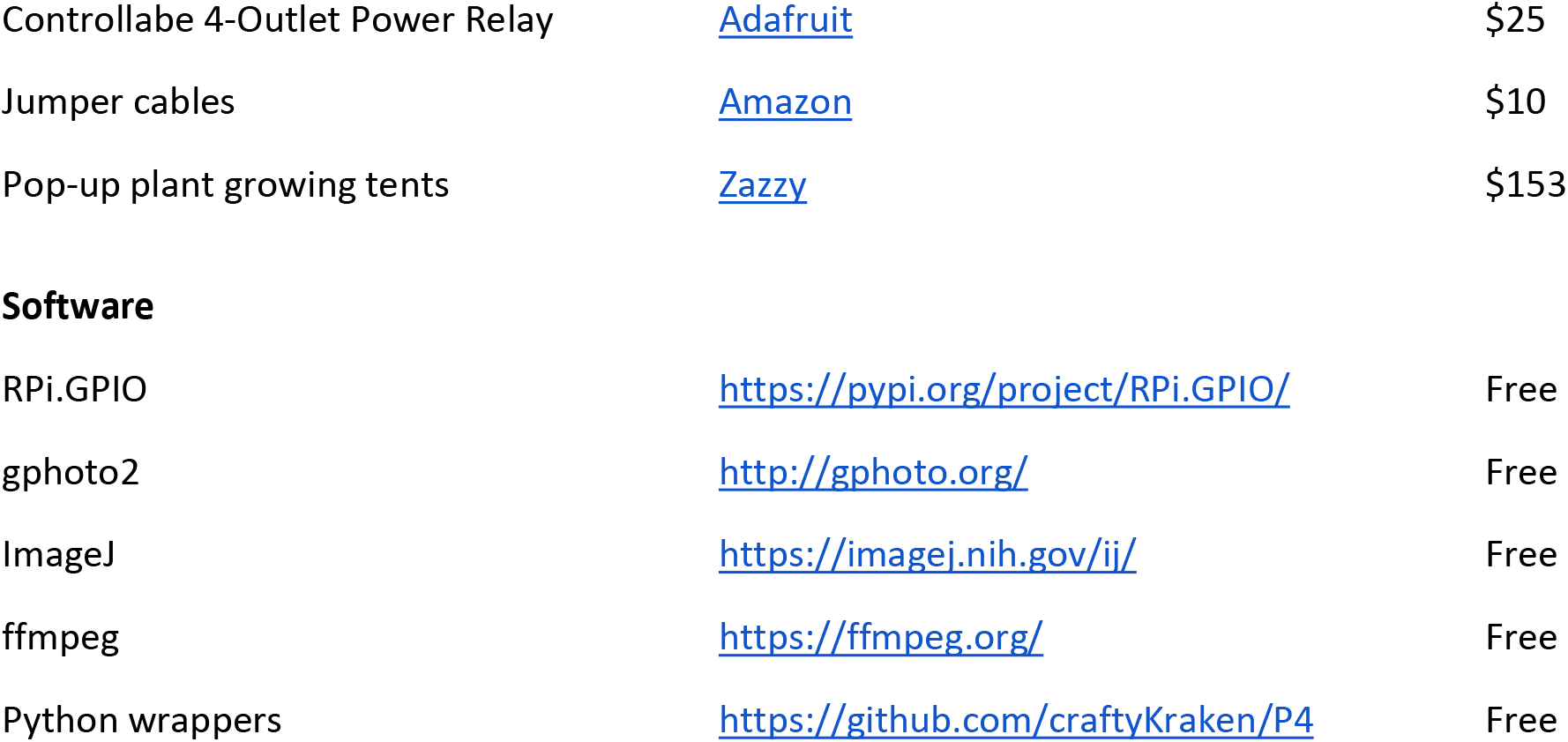
Components for DSLR-Based Imaging Platform

| **<u>Component</u>** | **<u>Source</u>** | **<u>Cost</u>** |
| --- | --- | --- |
| **Hardware** |  |  |
| RaspberryPi 3 Complete Starter Kit | CanaKit | \$75 |
| Canon EOS 80D DSLR | Canon | \$1,100 |
| Canon AC adapter and DC coupler | B&H Photo | \$150 |
| EF 100mm f/2.8 Macro Lens | Canon | \$600 |
| Tripod | Amazon | \$25 |
| Controllabe 4-Outlet Power Relay | Adafruit | \$25 |
| Jumper cables | Amazon | \$10 |
| Pop-up plant growing tents | Zazzy | \$153 |
| Software |  |  |
| RPi.GPIO | https://pypi.org/project/RPi.GPIO/ | Free |
| gphoto2 | http://gphoto.org/ | Free |
| ImageJ | https://imagej.nih.gov/ij/ | Free |
| ffmpeg | https://ffmpeg.org/ | Free |
| Python wrappers | https://github.com/craftyKraken/P4 | Free |

With programmatically controlled long exposures and a tripod stabilizer, the DSLR camera provides the best of both worlds: high resolution and detail for macroscopic plant subjects under multiple light conditions, and sufficient sensitivity to capture even low levels of bioluminescence signal. We do observe higher levels of thermodynamic noise than using cooled CCD camera setups, but this is easily filtered away using opensource software such as ImageJ^22^. An example for comparison is given in Supplementary Figure 3, where the same periwinkle petal infiltration was imaged under the CCD camera (Supplementary Figure 3A) and then the DSLR (Supplementary Figure 3B). The DSLR captured high resolution, true color images under both the light and dark conditions, while showing similar light sensitivity to the reporter.

In contrast to all-in-one tabletop chamber systems, our setup with a dark room or pop-up tent^23^ permits both much larger plant subjects and flexible positioning of the camera relative to the subject. For example, we took images of flower petal infiltrations on intact rose bushes a meter in size. These features make our platform a compelling alternative to commercially available systems at a fraction of the cost. We used this platform to capture high-resolution images of various plant subjects (Figure 2B, Figure 3D,E), and to perform time-course imaging for an FBP under an *AtRAB18* drought-inducible promoter (Figure 4D). These results illustrate how our platform, in conjunction with the FBP, enable low cost bioluminescence reporting for dynamic biological phenomena.

## Discussion

The results presented here demonstrate how the FBP can convert a common plant metabolite, caffeic acid, into a luciferin and turn it over to produce robust luminescence *in planta*, without the addition of any additional substrate. We go on to show how the FBP can be deployed as a useful addition to the bioluminescent reporter toolbox in plants. As it emits in the green spectrum^13^, it should be spectrally separable from firefly luciferase, allowing it to be applied in a complementary manner to the reporters that already exist. Our observation that the incorporation of the recycling pathway prolongs the auto-luminescent signal implies further tuning this pathway might be an avenue to enable long term imaging without substrate depletion in stable lines. While this may explain why we observe stronger auto-luminescence in infiltrated tissue than in the stable transgenic line, it might also be due to the overexpression of the enzymes due to the delivery of multiple T-DNAs. Thus, further optimizing the expression of the enzymes in the FBP might be another avenue to increase the auto-luminescent signal. This reengineering can be easily implemented in the future with our FBP toolkit, thanks to the easy promoter and terminator swapping enabled by using the MoClo system^14^.

Based on our transient expression assays, the FBP can function in a broad range of plant species. With the recent development of rapid morphogen mediated plant transgenesis protocols^24^, we believe FBP-based biosensors and reporters will be easy to extend across a range of plants for long term visualization of gene expression. This extensibility to a broad range of plants makes this approach to auto-luminescence superior to the existing transplastomic approach^11^, as plastid transformation is challenging and largely limited to tobacco and a few related *Solanaceae*^25^.

Our results with the pODO1 driven *Luz* in petunia flowers demonstrate that the FBP can be used to study the spatiotemporal patterns of gene expression, reporting the same patterns of gene expression as a firefly luciferase-based reporter^16^, but without necessitating substrate addition. The development of high expressing FBP lines by Mitiouchkina et al in parallel to this work demonstrate how our FBP tools could be applied to visualize gene expression in various tissues over the lifetime of the plant^29^. Thus, its application would lower the cost of performing these kinds of the experiments. Further, the data shows how by swapping promoters in the pathway, spatio-temporal patterns of auto-luminescence can be programmed into plants for synthetic biology applications. For example, plants engineered to have sufficiently bright auto-luminescent flowers might attract nocturnal insects enabling the creation of novel plant-pollinator interactions. The creation of bioluminescent flowers would also have relevance to designer floriculture and might serve as a visually compelling and viscerally exciting demonstration to the public of the power of synthetic biology. We plan to generate stable FBP petunia lines and explore this avenue of research further in the future.

The data we present on using the pRAB18 driven *Luz* to track drought stress via ABA signaling serves as a demonstration for how the FBP could be used to build biosensors for internal or external signals by driving genes in the pathway with synthetic promoters. In the future, stable lines generated with this construct could be deployed to track soil moisture levels at field scales. These biosensors would have an output more easily visualized than fluorescent reporters, have a high signal to noise ratio, and be cost effective due to their substrate independence. Thanks to the ease of promoter swapping in our FBP toolkit, this approach could be easily extended to build biosensors for a range of different phytohormones, environmental cues like light, or synthetic signaling systems for basic science and translational applications.

Besides the substrate costs, the high cost of commercially available bioluminescence imaging systems restrict access to luminescence-based reporters. We demonstrate how commercially available photography equipment and open source software can be used to set up relatively low-cost, modular and programmable luminescence imaging platforms. This coupled with the substrate independence of the FBP will broaden access to these tools to lower resource settings as well as enable scalable application of these reporters. We hope the work presented here will serve as a first step for the creation of novel bioluminescence-based tools in plants for basic science discovery and synthetic biology enabled applications.

## Materials and methods

### FBP plasmid construction

The various FBP encoding T-DNA constructs that were characterized in this work were built using a two-step process. First base plasmids containing an expression cassette was built for each of the five enzymes in the FBP. These base plasmids were designed to have promoters for either constitutive or tissue/time-period specific expression of the FBP using a mix of promoters that were either from the MoClo toolkit^14^ or amplified from genomic or plasmid DNA with primers that added the appropriate BsaI sites. The AtRAB18 and PhODO1 promoters were amplified from published plasmids that were obtained from either addgene or via requests from the authors. Terminators were chosen from the MoClo kit to ensure high expression^14^. DNA sequences encoding the enzymes were designed to be codon optimized for *N. benthamiana*. Additionally, all the common type-IIs restriction sites were removed via codon swaps. These sequences were synthesized by Twist biosciences. All these parts were assembled into the base vectors with a BsaI-based goldengate assembly reaction. The base vector backbones were designed to contain the appropriate AarI sites to be assembled together into either four or five expression cassettes containing T-DNAs through an AarI-based goldengate assembly reaction^26^. All vectors listed in Table 1 are available via addgene or upon request.

### Agrobacterium infiltration for transient expression of FBP

For all the transient expression assays of an FBP besides those shown in Figure 2A,B, the protocol described by Sparkes et al. was used^27^. Briefly, agrobacterium strains transformed with T-DNAs expressing the grown up overnight under Kanamycin and Gentamicin selection. These cultures were then pelleted and washed twice with infiltration media the next day and then resuspended at OD_600_ 0.8 and infiltrated into the desired tissue using a needleless syringe. Imaging of the luminescence signal was performed between three and five days after infiltration, depending on the assay being performed.

### AgroBEST co-culture for transient expression of FBP

The transient expression assays for *S. lycopersicum* and *A. thaliana* were carried out using the AgroBEST protocol^15^. Briefly, seeds were sterilized and germinated in 0.5x MS and once cotyledons emerged, the seedlings were co-cultured with the appropriate agrobacterium strain that had been transformed with the T-DNA encoding an FBP, in a mixture of MS and AB-MES. For all tomato AgroBEST treatments FBP_6 was used (Table 1). For the *Arabidopsis* AgroBEST FBP_12 and FBP_11 (Table 1) were used. These strains were primed for agrobest the previous day through overnight culture in AB-MES salts according to the AgroBEST protocol. Visualization of the bioluminescence signal was performed after two days of co-culture.

### Generation of a stable transgenic FBP line of *Nicotiana benthamiana*

The protocol detailed in Sparkes et al.^27^ was used to generate stable lines of *Nicotiana benthamiana* with the FBP integrated into the genome. To summarize, the agrobacterium strain characterized in Figure 1 was used to infiltrate leaves of *N. benthamiana*. After three days these leaves were excised, sterilized and transferred to shooting media plates that contained Kanamycin for selection. After callus and shoot formation, the shoots were transformed to rooting media plates which also had Kanamycin in the media. Finally, rooted plantlets were screened for luminescence and the individual with the highest signal was transferred to soil for seed.

### CCD camera-based luminescence imaging

For all static CCD camera-based luciferase imaging eight minute exposures of plant tissue were taken in using a UVP BioImaging Systems EpiChemi3 Darkroom. For transgenic plants a twelve minute exposure was used. Paired bright field images were also taken using the same camera. The brightness and contrast of the long exposure images were adjusted using imageJ^22^ to optemize signal visibility and then overlaid as a false colored image on the bright field image, where warmer colors correspond to higher signals.

### Time lapse CCD camera-based imaging

For the time lapse data collected in Figures 3A,C and in Figure 4C, experiments were performed using the NightOWL LB 983 *in vivo* imaging system^16^. In all cases agrobacterium infiltration of the FBP into the desired tissue was performed as described previously. For the petunia time lapse imaging data displayed in Figure 3A and C, the petals of flowers were infiltrated and then the flowers were excised from the plants either one or two days after infiltrations and were mounted in the imaging platform with their pedicles immersed in a solution of 5% sucrose^16^. Images were then captured every hour with a ten-minute exposure. Long day light conditions were implemented in the times between images. For the data displayed in Figure 4C, leaves of *N. benthamiana* were infiltrated and then excised from the plant after three days and placed in petri dishes. For the watered conditions a moistened filter paper was added to the petri dish, whereas for the drought conditions the filter paper was left dry. Images were then captured every hour with a ten-minute exposure and the timepoint with the peak pRAB18:Luz signal was reported.

### Quantification of luminescence signal in images

For quantification of luminescence signal, imageJ^22^ was used to box areas of the same size in images of infiltrated leaves or 96 well plates containing hole punches from infiltrated leaves and then the average signal intensity was recorded and plotted in python using the seaborn^28^ plotting package. All p values reported were calculated using the t-test function in the scipy package. All the raw data and data analysis code were made available on Github (https://github.com/craftyKraken/P4).

### Time lapse luminescence imaging using DSLR-based system

A complete listing of sourced components and costs is found in Table 2. Timelapse images were captured using a Canon DSLR equipped with a macro lens and programmably controlled from a Raspberry Pi microcomputer running a combination of free, open-source and custom software. The gphoto2 library was used to control the camera via USB; power to lights and the camera was modulated by a pair of controllable four outlet power relays connected to the Pi GPIO interface via jumper cables; a custom python wrapper was used to implement control logic for the camera, using gphoto2, and for the relay, using the RPi.GPIO package. Long-exposure images were captured in a dark room or light-proof pop-up tent. Image processing was performed using macros written in ImageJ, and the FFMPEG library, under top-level control of an additional custom python wrapper. Gphoto2, ImageJ and FFMPEG are free and open-source software; the custom wrappers are hosted on a public GitHub repository (https://github.com/craftyKraken/P4), and are also available for reuse under the GNU 3.0 public license.

---

**Supplementary Figure 1.**
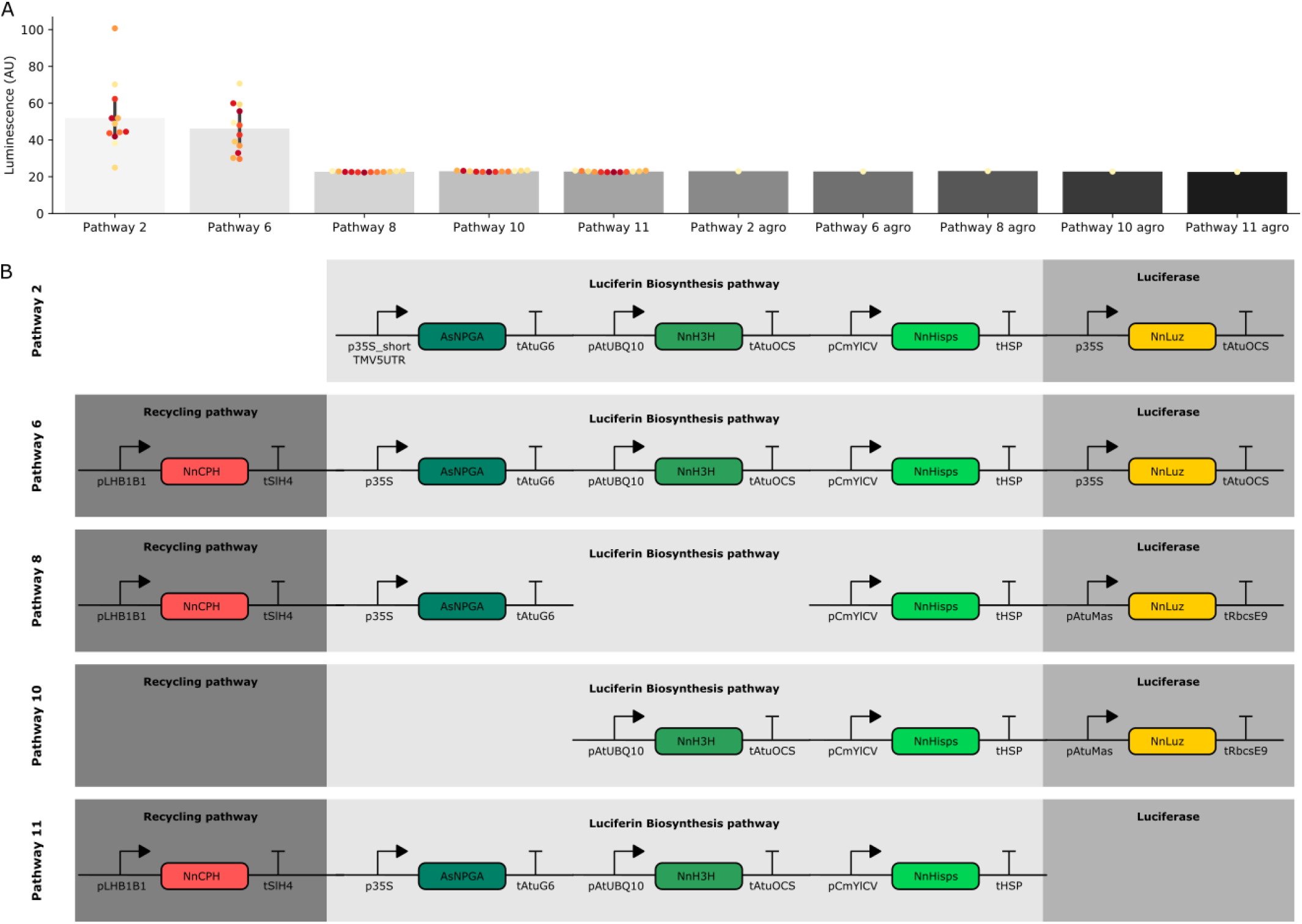
A) Bar plots summarizing luminescence recorded from *N. benthamiana* leaves infiltrated with FBPs three days after infiltration. Each dot represents a reading from a different infiltrated leaf. The back bars represent standard deviations. The background level of Luminescence recorded was 23 AU. The bars labeled agro are readings from confluent cultures of agrobacterium strains used to deliver the respective FBP. Pathway 2 is a functional pathway without CPH and pathway 6 is a functional pathway with CPH. Pathways 8,10 and 11 are all negative controls missing pathway enzymes. B) Schematics of the expression cassettes assembled for each of the pathways characterized.

**Supplementary Figure 2.**
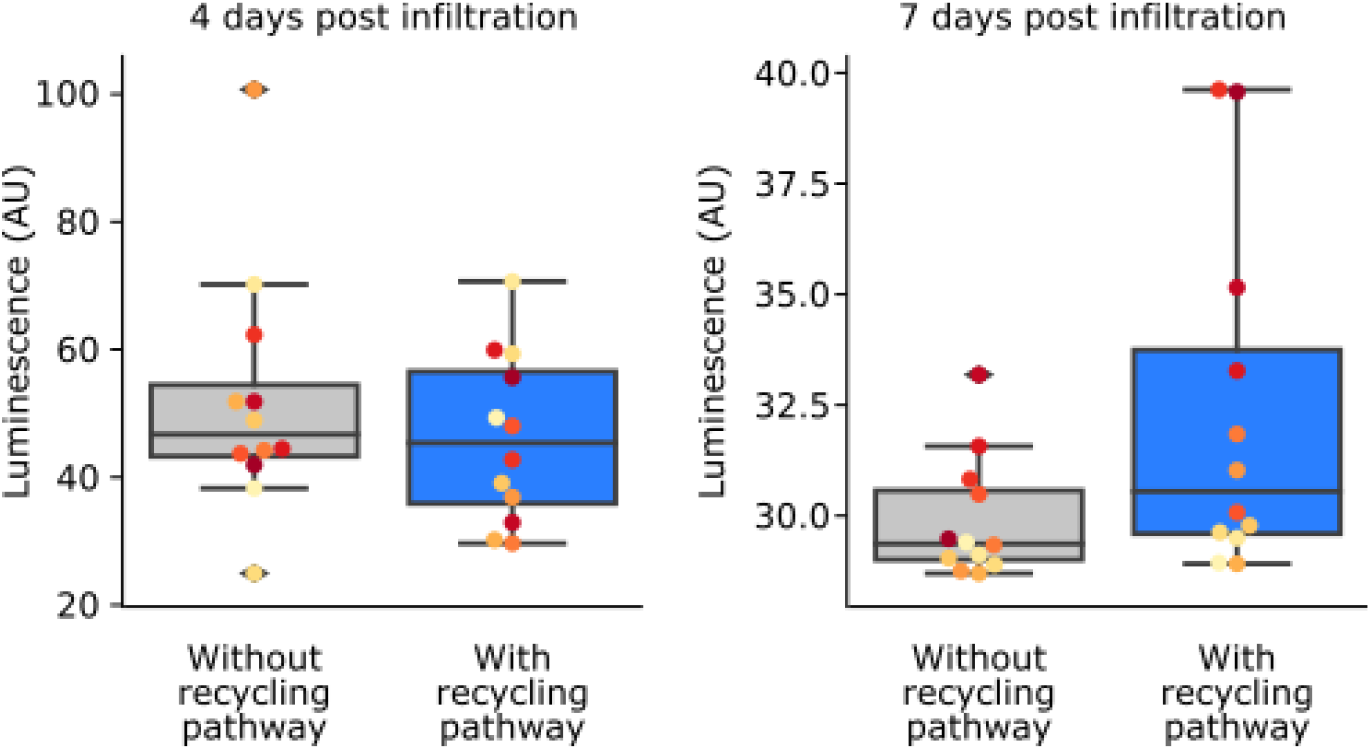
Box plots visualizing luminescence from *N. benthamiana* leaves infiltrated an FBP that either with (blue) or without (gray) the *CPH*-based recycling pathway. Each dot represents a reading from a different infiltrated leaf. Data is form four days and seven days after infiltration.

**Supplementary figure 3.**
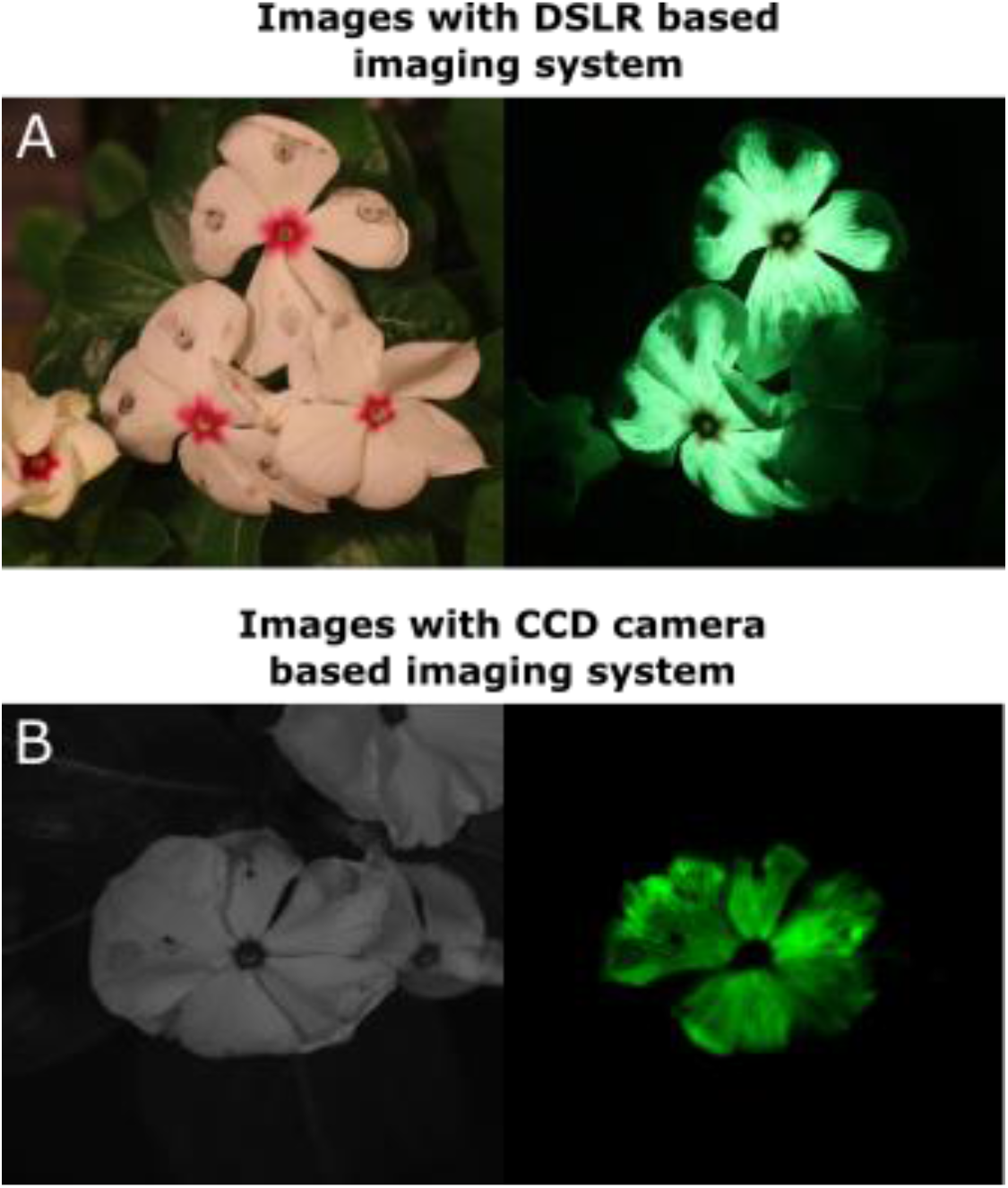
Petal infiltrations of *C. roseus* flowers with an FBP imaged three days post infiltration with either our DLSR-based imaging system (A) or a CCD based imaging system (B).

**Video figure 1**. Time lapse imaging was performed with our RaspberryPi controlled DSLR imaging platform to visualize ABA induced FBP expression in plants exposed to drought stress. We infiltrated one leaf each on two *N. benthamiana* plants with FBPs that had either a 35S (+control) or pRAB18 driven *Luz*. One plant was allowed to desiccate while the other was kept watered. Bioluminescence in the leaves as then imaged over three days. The treatments of the leaves are labeled with white legends.

## Acknowledgements

N.L. and T.I. are supported by grants from National Institute of Health (R01GM079712) and Next-Generation BioGreen 21 Program (PJ013386, Rural Development Administration, Republic of Korea). We would like to thank Dr. Karen Sarkisyan and Dr. Nadya Markina for kindly sharing the sequences of the FBP genes from their publication.

## References

1. Shimomura, O. (2006). Bioluminescence: chemical principles and methods. World Scientific Publishing Co. Pte.

2. Wainwright, P. C., & Longo, S. J. (2017). Functional innovations and the conquest of the oceans by Acanthomorph fishes. Current Biology, 27(11), R550–R557.

3. Verdes, A., & Gruber, D. F. (2017). Glowing worms: Biological, chemical, and functional diversity of bioluminescent annelids. Integrative and comparative biology, 57(1), 18–32.

4. Labella, A. M., Arahal, D. R., Castro, D., Lemos, M. L., & Borrego, J. J. (2017). Revisiting the genus Photobacterium: taxonomy, ecology and pathogenesis. Int Microbiol, 20, 1–10.

5. Oliveira, A. G., Stevani, C. V., Waldenmaier, H. E., Viviani, V., Emerson, J. M., Loros, J. J., & Dunlap, J. C. (2015). Circadian control sheds light on fungal bioluminescence. Current Biology, 25(7), 964–968.

6. Oba, Y., Stevani, C. V., Oliveira, A. G., Tsarkova, A. S., Chepurnykh, T. V., & Yampolsky, I. V. (2017). Selected least studied but not forgotten bioluminescent systems. Photochemistry and photobiology, 93(2), 405–415.

7. Schultz, D. T., Kotlobay, A. A., Ziganshin, R., Bannikov, A., Markina, N. M., Chepurnyh, T. V., … & Oba, Y. (2018). Luciferase of the Japanese syllid polychaete Odontosyllis umdecimdonta. Biochemical and biophysical research communications, 502(3), 318–323.

8. Khakhar, A., Leydon, A. R., Lemmex, A. C., Klavins, E., & Nemhauser, J. L. (2018). Synthetic hormone-responsive transcription factors can monitor and re-program plant development. Elife, 7, e34702.

9. Wend, S., Dal Bosco, C., Kämpf, M. M., Ren, F., Palme, K., Weber, W., … & Zurbriggen, M. D. (2013). A quantitative ratiometric sensor for time-resolved analysis of auxin dynamics. Scientific reports, 3, 2052.

10. Rellán-Álvarez, R., Lobet, G., Lindner, H., Pradier, P. L., Sebastian, J., Yee, M. C., … & Haney, C. H. (2015). GLO-Roots: an imaging platform enabling multidimensional characterization of soil-grown root systems. Elife, 4, e07597.

11. Krichevsky, A., Meyers, B., Vainstein, A., Maliga, P., & Citovsky, V. (2010). Autoluminescent plants. PloS one, 5(11), e15461.

12. Kotlobay, A. A., Sarkisyan, K. S., Mokrushina, Y. A., Marcet-Houben, M., Serebrovskaya, E. O., Markina, N. M., … & Petushkov, V. N. (2018). Genetically encodable bioluminescent system from fungi. Proceedings of the National Academy of Sciences, 115(50), 12728–12732.

13. Kaskova, Z. M., Dörr, F. A., Petushkov, V. N., Purtov, K. V., Tsarkova, A. S., Rodionova, N. S., … & Baranov, M. S. (2017). Mechanism and color modulation of fungal bioluminescence. Science advances, 3(4), e1602847.

14. Engler, C., Youles, M., Gruetzner, R., Ehnert, T. M., Werner, S., Jones, J. D., … & Marillonnet, S. (2014). A golden gate modular cloning toolbox for plants. ACS synthetic biology, 3(11), 839–843.

15. Wu, H. Y., Liu, K. H., Wang, Y. C., Wu, J. F., Chiu, W. L., Chen, C. Y., … & Lai, E. M. (2014). AGROBEST: an efficient Agrobacterium-mediated transient expression method for versatile gene function analyses in Arabidopsis seedlings. Plant methods, 10(1), 19.

16. Fenske, M. P., Hazelton, K. D. H., Hempton, A. K., Shim, J. S., Yamamoto, B. M., Riffell, J. A., & Imaizumi, T. (2015). Circadian clock gene LATE ELONGATED HYPOCOTYL directly regulates the timing of floral scent emission in Petunia. Proceedings of the National Academy of Sciences, 112(31), 9775–9780.

17. Van Moerkercke, A., Haring, M. A., & Schuurink, R. C. (2011). The transcription factor EMISSION OF BENZENOIDS II activates the MYB ODORANT1 promoter at a MYB binding site specific for fragrant petunias. The Plant Journal, 67(5), 917–928.

18. Verdonk, J. C., Haring, M. A., van Tunen, A. J., & Schuurink, R. C. (2005). ODORANT1 regulates fragrance biosynthesis in petunia flowers. The Plant Cell, 17(5), 1612–1624.

19. Norkunas, K., Harding, R., Dale, J., & Dugdale, B. (2018). Improving agroinfiltration-based transient gene expression in Nicotiana benthamiana. Plant methods, 14(1), 71.

20. Kim, T. H., Hauser, F., Ha, T., Xue, S., Böhmer, M., Nishimura, N., … & Lee, S. (2011). Chemical genetics reveals negative regulation of abscisic acid signaling by a plant immune response pathway. Current Biology, 21(11), 990–997.

21. Reddy, A. R., Chaitanya, K. V., & Vivekanandan, M. (2004). Drought-induced responses of photosynthesis and antioxidant metabolism in higher plants. Journal of plant physiology, 161(11), 1189–1202.

22. Schindelin, J., Arganda-Carreras, I., Frise, E., Kaynig, V., Longair, M., Pietzsch, T., … & Tinevez, J. Y. (2012). Fiji: an open-source platform for biological-image analysis. Nature methods, 9(7), 676.

23. Katagiri, F., Canelon-Suarez, D., Griffin, K., Petersen, J., Meyer, R. K., Siegle, M., & Mase, K. (2015). Design and construction of an inexpensive homemade plant growth chamber. PloS one, 10(5), e0126826.

24. Lowe, K., La Rota, M., Hoerster, G., Hastings, C., Wang, N., Chamberlin, M., … & Gordon-Kamm, W. (2018). Rapid genotype “independent” Zea mays L.(maize) transformation via direct somatic embryogenesis. In Vitro Cellular & Developmental Biology-Plant, 54(3), 240–252.

25. Rigano, M. M., Scotti, N., & Cardi, T. (2012). Unsolved problems in plastid transformation. Bioengineered, 3(6), 329–333.

26. Čermák, T., Curtin, S. J., Gil-Humanes, J., Čegan, R., Kono, T. J., Konečná, E., … & Voytas, D. F. (2017). A multipurpose toolkit to enable advanced genome engineering in plants. The Plant Cell, 29(6), 1196–1217.

27. Sparkes, I. A., Runions, J., Kearns, A., & Hawes, C. (2006). Rapid, transient expression of fluorescent fusion proteins in tobacco plants and generation of stably transformed plants. Nature protocols, 1(4), 2019.

28. Waskom, M., Botvinnik, O., Hobson, P., Cole, J. B., Halchenko, Y., Hoyer, S., … & Coelho, L. P. (2014). seaborn: v0. 5.0 (November 2014). Zenodo, doi, 10.

29. Tatiana Mitiouchkina, Alexander S. Mishin, Louisa Gonzalez Somermeyer, Nadezhda M. Markina,Tatiana V. Chepurnyh, Elena B. Guglya, Tatiana A. Karataeva, Kseniia A. Palkina, Ekaterina S. Shakhova, Liliia I. Fakhranurova, Sofia V. Chekova, Aleksandra S. Tsarkova, Yaroslav V. Golubev, Vadim V. Negrebetsky, Sergey A. Dolgushin, Pavel V. Shalaev, Olesya A. Melnik, Victoria O. Shipunova, Sergey M. Deyev, Andrey I. Bubyrev, Alexander S. Pushin, Vladimir V. Choob, Sergey V. Dolgov, Fyodor A. Kondrashov, Ilia V. Yampolsky and Karen S. Sarkisyan. “Plants with self-sustained luminescence”, 2019 Biorxiv

